# Impact of a standardized *Tetradesmus obliquus* Mi175.B1.a extract on gut function and physiology in inflammatory conditions: *in vitro* and *in vivo* insights

**DOI:** 10.1101/2025.05.21.655247

**Authors:** Jonathan Maury, Pierre Albina, Anne Abot, Rémi Pradelles

## Abstract

Microalgae have emerged as a promising source of natural nutraceuticals, which may play a critical role in maintaining the structural integrity of the gut mucosal barrier and modulating immune responses. Extracts from *Tetradesmus obliquus* have demonstrated potential benefits for gut health in humans. This study aimed to evaluate the efficacy of *Tetradesmus obliquus* Mi175.B1.a through two complementary approaches: (1) an *in vitro* model using the HT-29 cell line to assess cytotoxicity, intestinal permeability, immunomodulatory properties, and gut barrier function, and (2) an *in vivo* mouse model of dextran sodium sulfate (DSS)-induced colitis to examine parameters relevant to digestive physiology. In the *in vitro* model, HT-29 cells were pretreated with two selected doses of *T. obliquus* (Mi175.B1.a) extract for 72 hours prior to stress induction with lipopolysaccharide (LPS). Parameters assessed included gene expression, paracellular permeability, and immunomodulatory properties. For the *in vivo* model, forty (40) male mice were randomly allocated to receive daily oral administration of either vehicle or *T. obliquus* extract at two different doses for 21 days. Colitis was induced from Day 14 to Day 21 using 3% DSS, and macroscopic and gut motility parameters were subsequently evaluated. Key findings revealed that *T. obliquus* (Mi175.B1.a) significantly reduced FITC-dextran permeability, restored physiological expression of *Zonula occludens-1* (a tight junction protein), and attenuated TNF-α release and *Cox2* gene expression in comparison to the LPS-only condition. In the *in vivo* model, *T. obliquus* extract significantly reduced the daily disease activity index and colitis score compared to DSS-only condition. Importantly, the two evaluated doses of *T. obliquus* (Mi175.B1.a) were well-tolerated, demonstrating no toxic effects. These results highlight the potential of *T. obliquus* extract to enhance intestinal barrier integrity by regulating inflammatory and oxidative stress pathways opening promising research perspectives to determine the benefits in human application.

## 1. Introduction

The intestinal barrier, sustained by the gut epithelium, microbiome, and immune system, depends on the integrity of junctional proteins, including tight junctions, desmosomes, and adherens junctions, to maintain homeostasis and prevent the translocation of luminal substances and pathogens into the internal environment [1]. This barrier is a dynamic system influenced by the intestinal microbiome composition and intercellular connection activity, which are regulated by hormones, dietary components, inflammatory mediators, and the enteric nervous system [2]. Increased gut permeability, commonly referred to as “leaky gut,” results from disruptions in the structural and functional integrity of the intestinal barrier. Beyond environmental factors, key contributors to this condition include bacterial infections, inflammation, oxidative stress, and microbial dysbiosis. Dysbiosis, characterized by an imbalance in the composition, diversity, or functional activities of the gut microbiome, is particularly significant. It is associated with reduced production of protective short-chain fatty acids (SCFAs) and dysregulation of tight junction proteins such as zonula occludens-1 and occludin [1,3–6]. Dysbiosis can disrupt intestinal homeostasis by promoting the overexpression of pro-inflammatory cytokines, the overgrowth of Gram-negative, lipopolysaccharide-producing Proteobacteria, and a decline in microbial richness and diversity [4–6]. Impairment of the intestinal barrier can lead to a spectrum of health consequences, ranging from acute gastrointestinal disorders to chronic pathological conditions, including metabolic diseases, inflammatory bowel disease (IBD), irritable bowel syndrome (IBS), neurodegenerative disorders, and others [3]. Given the high prevalence of gastrointestinal (GI) disorders and symptoms, which impact over 40% of the global population, identifying effective therapeutics and preventive approaches to improve the gut microbiome and maintain intestinal barrier integrity is of critical importance. In this regard, dietary interventions represent a particularly promising approach to restore gut microbiota balance [7].

Beyond conventional prebiotics, probiotics, and postbiotics, a growing body of research has identified additional nutraceuticals, particularly plant-derived extracts, as potential modulators of gut health [8]. Phytochemicals have garnered significant interest due to their prebiotic-like effects, attributed to their antioxidant and anti-inflammatory properties, as well as their limited absorption, which prolongs their retention time in the intestine, where they may positively influence gut barrier integrity and microbiota composition [9,10]. Furthermore, the nutraceutical industry is evolving rapidly, driven by increasing demand for natural and sustainable products in response to ecological concerns and biodiversity conservation. In this context, microalgae-based ingredients represent a promising alternative, providing a rich source of bioactive compounds, including essential nutrients (e.g., vitamins, minerals, polyunsaturated fatty acids) and phytochemicals such as pigmented compounds, polyphenols, and sterols, which are well recognized for their involvement in key pathways regulating gut function, microbiota composition, and intestinal barrier integrity [11–13]. For instance, extracts from *Spirulina platensis, Dunaliella salina*, and *Chlorella vulgaris* have demonstrated potential in alleviating gastrointestinal disorders by reducing mucosal damage, inflammation, and oxidative stress in colon tissue, while modulating gut microbiota in various murine models [14–16]. Among microalgae species, *Tetradesmus obliquus* has been widely recognized for its suitability in large-scale biomass production, particularly using photobioreactor technology, due to its rapid resistance to adverse environmental conditions [17]. This microalgal strain contains multiple bioactive molecules, including essential fatty acids (e.g., α-linolenic acid), carotenoids (e.g., lutein), carbohydrates, and proteins (e.g., leucine/isoleucine), which have been suggested to enhance microbiota diversity and reinforce gut barrier function [18–24]. Recent findings indicate that various microalgae supplementation, including *T. obliquus* dry biomass, may support gastrointestinal health in dogs by selectively stimulating beneficial microbiota without compromising food intake or digestibility [25]. However, further studies in specific models are necessary to validate these results and explore the potential application of *T. obliquus*-derived ingredients for human gut health *in vivo* as complementary approach to evaluate the impact on gut.

The use of an *in vitro* (e.g. cell cultures, isolated tissue) and *in vivo* (e.g. mice models) are complementary approaches to study underlying mechanism of actions, capture the full physiological impact including metabolic and systemic effects and identify dose(s) of efficacy. In particular in pro-inflammatory conditions, *in vitro* and *in vivo* models have been developed to study numerous functions of the intestinal barrier and gut microbiota with a demonstrated sensitivity to nutraceutical applications [26–33]. Among intestinal epithelial cells, the human colon cell line HT-29 is well recognized to study human colon cancer but this cell-line model provide a reproducible and human-relevant approach to assess intestinal permeability and immune interactions [28]. These cells, derived from human colorectal adenocarcinoma, exhibit key features of the intestinal epithelium, including the formation of a polarized monolayer and the expression of tight junction proteins. This makes them a valuable model for studying the effects of bioactive compounds on gut barrier integrity. This cell-based assay is particularly relevant for assessing the efficacy of dietary supplements, probiotics, or pharmaceutical compounds aimed at improving digestive health and reducing intestinal inflammation. A recent study showed benefits of a combination of several ingredients (e.g. L-glutamine, vitamins, grape-seed extract etc.) on different parameters evaluating gut barrier integrity such as transfer FITC-dextran, mRNA expression of thigh junction proteins, cytokine levels in apical compartment and antioxidant capacities in pro-inflammatory conditions [28]. For *in vivo* model, Dextran Sulphate Sodium (DSS)-induced colitis in mice is widely used to mimic the negative impact of inflammatory bowel disease by inducing intestinal epithelial barrier damage and increasing gut permeability [34]. DSS-induced colitis model works by the activation of immune cells then triggers characterized the production of inflammatory cytokines, an increase in reactive oxygen species and thus oxidative stress, and by destabilizing inter-cellular junctions due to internalization and degradation of junction proteins [35–38]. Beyond biological parameters, it’s also interesting to evaluate in DSS-induced colitis the impact on global symptoms such as daily disease activity (DAI), colitis or colon amplitude scores [30,32,39,40].

Thus, the aim of this study was to evaluate the efficacy of *Tetradesmus obliquus* Mi175.B1.a extract in 1/ an *in vitro* test using HT-29 cell line on cytotoxicity, intestinal permeability, immunomodulatory properties and gut barrier function and 2/ in a dextran sodium sulfate (DSS) model of mouse colitis on several parameters relevant to digestive physiology.

## 2. Results

### 2.1. Effect on in vitro model parameters

#### 2.1.1. Cytotoxicity analysis

Before evaluating the effects of test items on Intestinal Epithelial Cells (IEC), it was necessary to evaluate the cytotoxicity of test items to choose test items concentrations for next steps. Tested items were added in the culture media at the indicated concentrations for 24h, 48h and 72h of treatment. The cytotoxicity was evaluated by the lactate dehydrogenase (LDH) activity measurement and the viability was evaluated by the formazan production measurement. The vehicle tests for each dilution were performed with the corresponding concentration of EtOH (i.e.: vehicle D1 = 0.1% of each, vehicle D2 = 0.02% of each, vehicle D3 = 0.01% of each, vehicle D4 =0.002% of each and vehicle D5 =0.0002% of each). For each condition, morphological aspect of cells was appreciated after the experimentation, and we did not observe any morphological modifications of the cells compared to the control condition. As shown in Figure 1, we did not observe cytotoxicity or loss of cell viability in response to *T. obliquus* Mi.175.B1.a extract, regardless of the dose used. For the next steps, we used the working concentrations of microalgae extract 10 (Dose 1 – D1) and 50 (Dose 2 – D2) mg/L.

**Figure 1.**
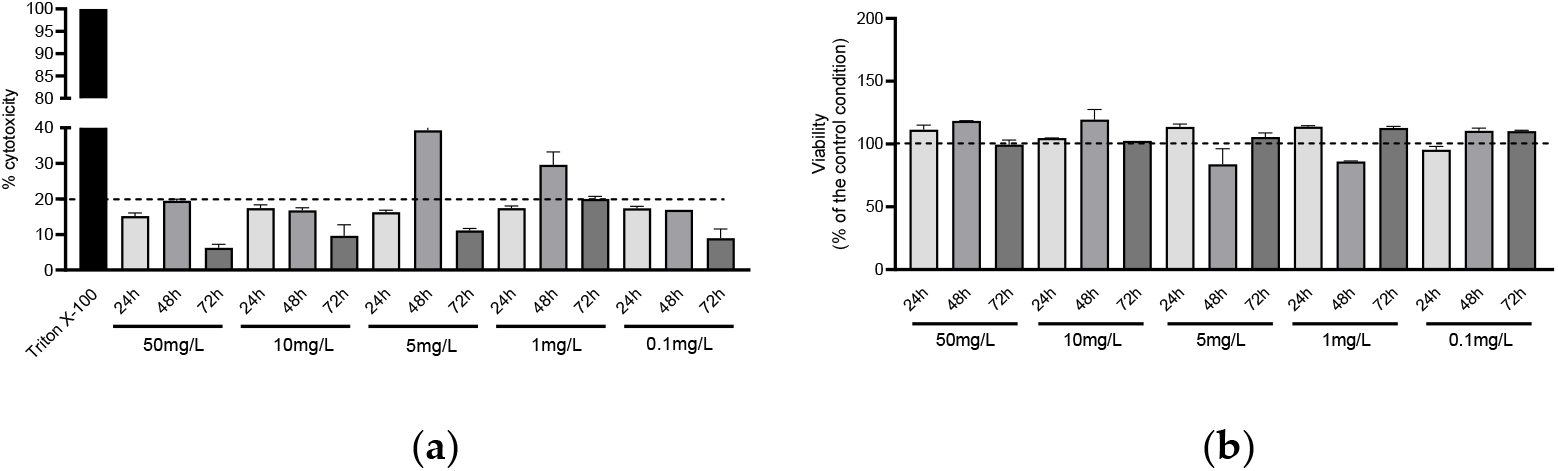
a) Evaluation of LDH released by IEC and b) Evaluation formazan production by IEC in response to *T. obliquus* Mi175.B1.a extract (n=2/group)

#### 2.1.2. Intestinal permeability and barrier function assessment

The intestinal permeability was assessed using a fluorometric assay and the results are presented in Figure 2. The FITC-dextran concentration from the basolateral side was correlated with the severity of permeability alteration of IEC. In a pro-inflammatory environment, induced by LPS, we observed an increase in the transfer of dextran in basolateral side that reflected the worsened integrity of IEC tight junctions. Microalgae treatments were able to reverse this deleterious effect induced by LPS regardless the tested concentration (p < 0.0001; F=3125,5; η^2^=0,99, large effect size). These results were correlated with the significant restoration of the expression of tight junctions *Zonula Occludens-1* (p = 0,01; F=27,56; η^2^=0,95, large effect size) and while only a trend was described for Claudin 4 (p = 0.081 ; F=4,840; η^2^=0,78, large effect size). Then, we studied the gene expression of biomarkers involved in intestinal homeostasis. The *Muc-2* gene coding for the oligomeric mucus gel-forming Mucin 2 provides an insoluble mucous barrier that serves to protect the colonic epithelium. Its expression was not significantly changed in all experimental groups (Figure 2.d.)

**Figure 2.**
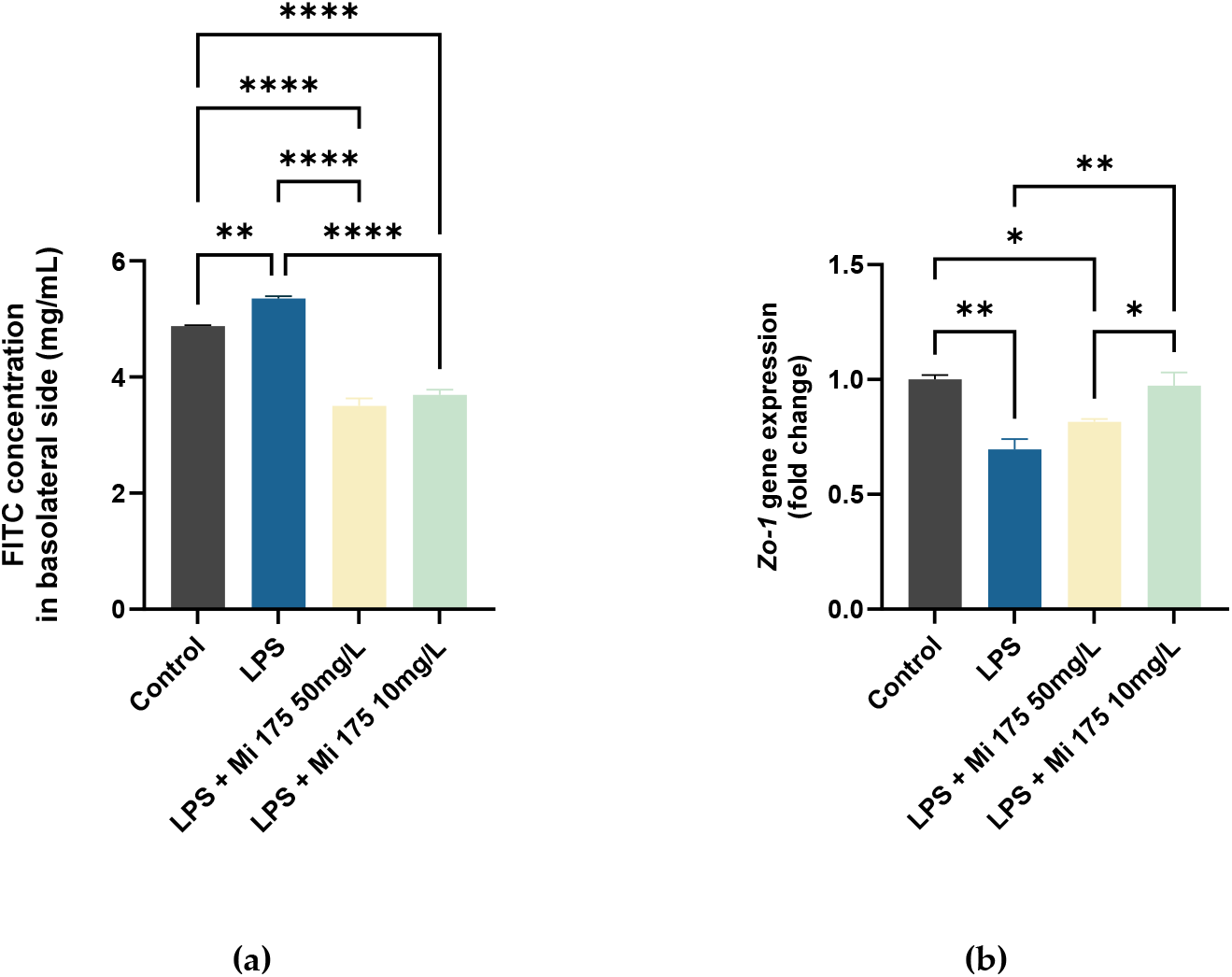
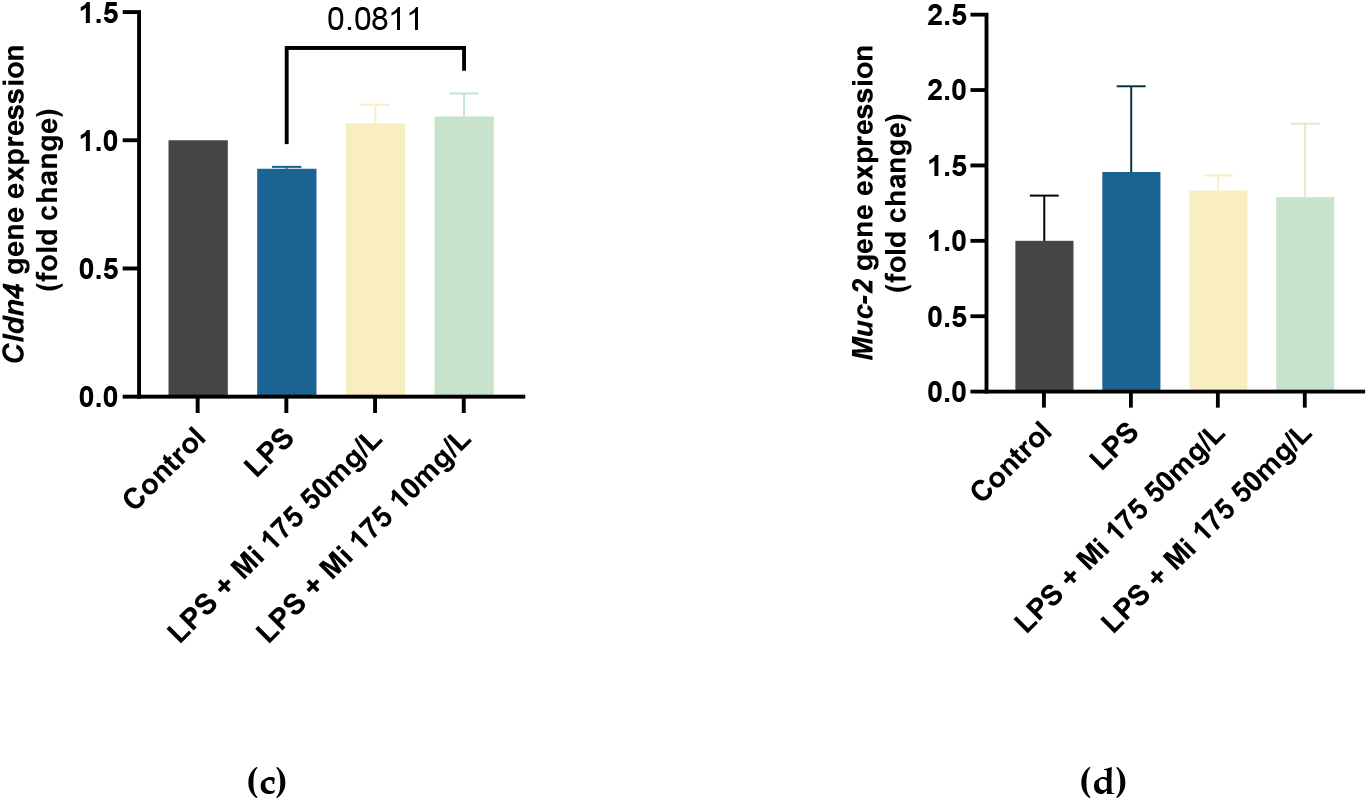
Intestinal permeability and barrier function properties of *T. obliuus* Mi175.B1.a extract in IEC. (a) permeability of IEC monolayer in response to exposure to LPS with or without microalgal solutions measured by transfer of fluorescent dextran, (b) Zo-1 expression was quantified using RT-qPCR in IEC, (c) Cldn4 expression was quantified using RT-qPCR in IEC (d) Muc-2 expression was quantified using RT-qPCR in IEC. * indicates significant difference between conditions using one-way ANOVA followed by Tukey’s post-hoc test. * means a p<0,05; ** means a p<0,01, *** means a p<0,001 and **** means a p<0,0001. n=2 / group.

#### 2.1.3. Immune modulation capacities assessment

The quantification of the expression of pro-inflammatory and anti-inflammatory cytokines was performed in response to LPS with or without the solutions of microalgae extract and the results are presented in Table 1.

**Table 1.**
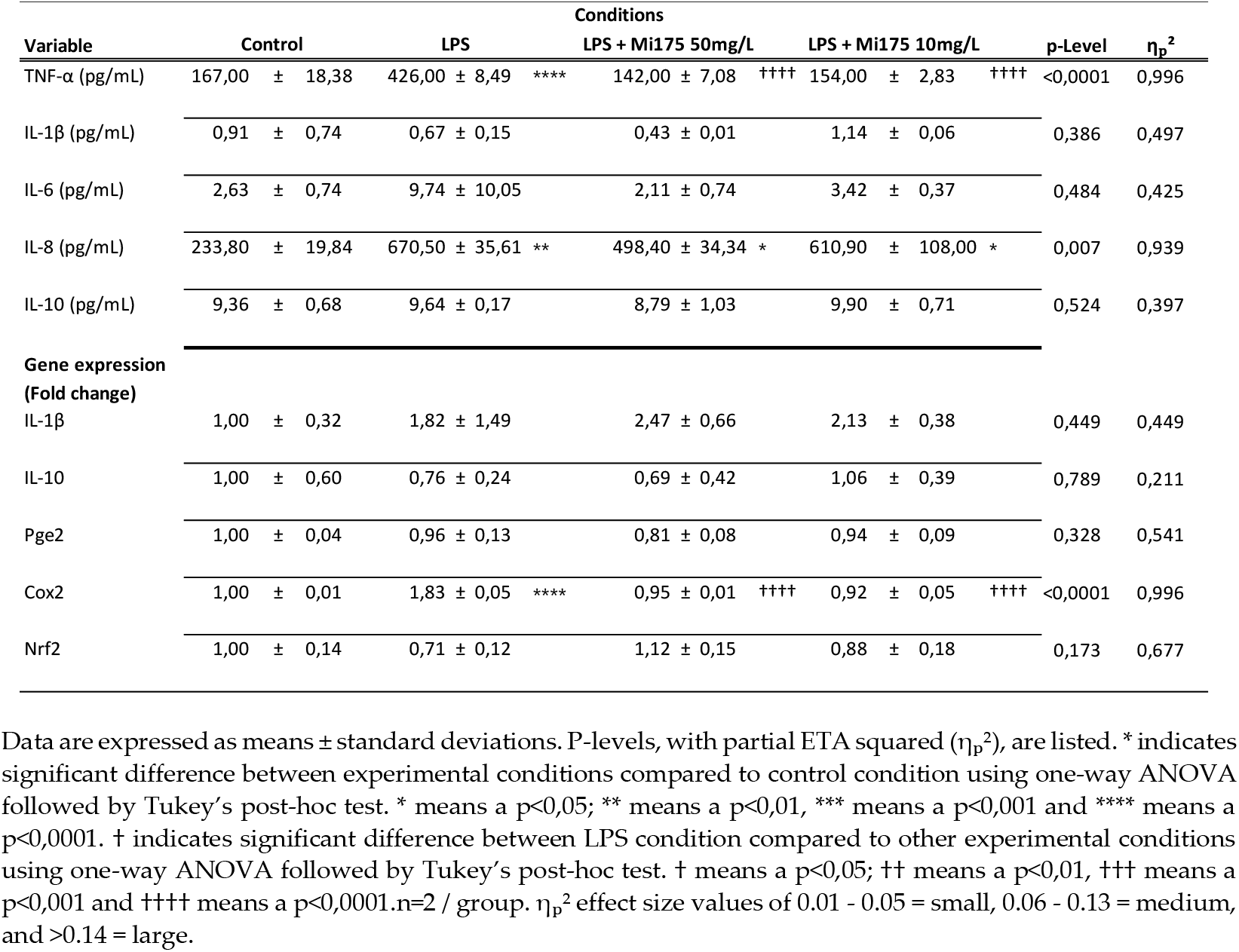
Immunomodulation properties, inflammatory response and oxidative stress defense biomarkers.

Data showed that LPS induced-stress strongly enhanced pro-inflammatory cytokine IL-8 and TNFα production. The induction of TNFα secretion in response to LPS was significantly blocked by *T. obliquus* (Mi175.B1.a) treatments, regardless of the concentration used with a large effect size. Concerning the other proinflammatory cytokines assessed, we did not observe significant variations of either IL-6 or IL-1β secreted in the culture media.

Then, we evaluated the expression of biomarker involved in regulation of intestinal homeostasis. The cyclooxygenase enzyme Cox2 is known to be induced during inflammatory conditions in the gut and play a key role in gut-related inflammatory disorders. We observed an enhanced expression of Cox2 after LPS stimulation. Interestingly microalgae treatments were able to fully reverse this effect regardless of the concentration tested with a large effect size. In our experimental condition, we did not observe any significant variation in the expression of other gene such as Prostaglandin E2 (PGE2) or Nrf2.

### 2.2. Effect on in vivo model parameters

#### 2.2.1. Metabolic parameters and animal observations assessment

First, mice were randomly assigned to the experimental groups without difference of body weight between the groups at D0. The body weight was assessed weekly (on Monday, 9:00 a.m.) during microalgae extract supplementation and then daily when DSS was added in the tap water. In comparison with the Vehicle-treated group (non-colitic group), the administration of DSS decreased body weight with significant group (p < 0.01 F(3; 36)=4,539; η2=0,266), time (p < 0.0001 ; F(1,124; 30,48)=233,0; η2=0,895) and Group*Time interaction (p < 0.0001 ; F(27; 324)=41,66; η2=0,776) effects (Figure 3.a.). Posthoc pairwise analysis revealed no significant difference between DSS-colitis and the two microalgae groups. The daily oral administration of *T. obliquus* (Mi175.B1.a) treatments in a preventive manner to mice did not show any sign of toxicity, which was evaluated by body weight increase, food intake, and general appearance of the animals. The addition of 3% (w/v) of DSS to drinking water for 6 days resulted in a progressive increase in daily Disease Activity Index (DAI) values in the DSS-colitis and the two microalgae groups, due to body weight loss and excretion of diarrheic/bleeding feces (Figure 1D). 2-way ANOVA analysis showed significant group (p < 0.0001 ; F(3; 36)=81,55; η2=0,871), time (p < 0.0001 ; F(3,860; 139)=233,0; η2=0,866) and Group*Time interaction (p < 0.0001 ; F(21; 252)=23,64; η2=0,663) effects. Post-hoc pairwise analysis interestingly showed the oral microalgae-treatment with 100mg/L significantly attenuated the impact of the DSS damage at Days 6 and 7 (Figure 1.b.).

**Figure 3.**
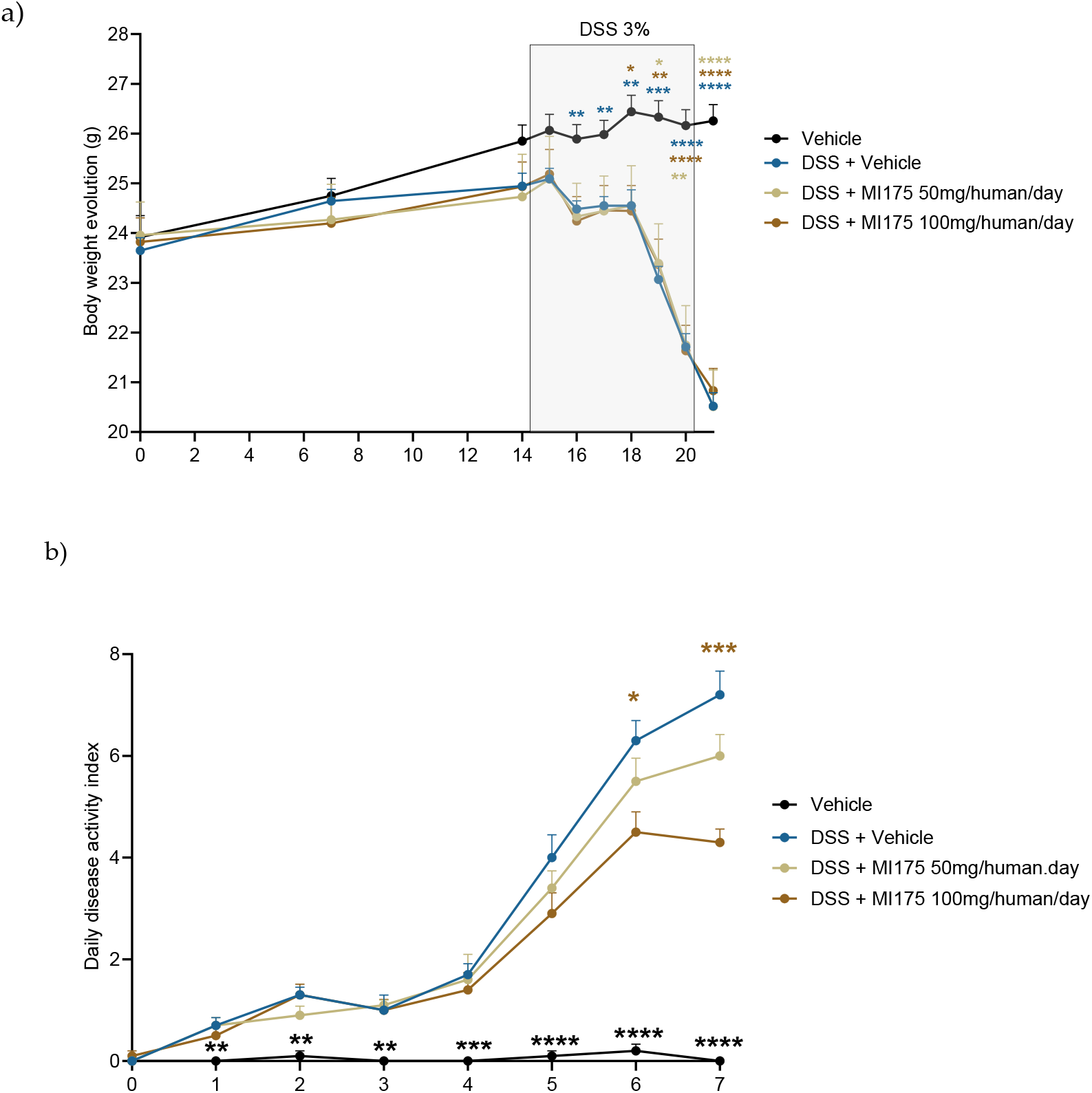
DSS-induced colitis monitoring. Effects of an oral administration of vehicle or 2 different doses of microalgal strain *T. obliquus* Mi175.B1.a extract on (a) body weight over time and (b) disease activity index (DAI) values evolution overtime. * indicates significant difference between conditions using 2-way ANOVA followed by Tukey’s post-hoc test. * means a p<0,05; ** means a p<0,01, *** means a p<0,001 and **** means a p<0,0001. n=9-10/ group.

#### 2.2.2. Macroscopic scoring assessment

The macroscopic evaluation of the colonic segments confirmed the beneficial effects found in microalgal extract-treated colitis mice. We showed a significant reduction in colonic weight/length ratio compared to the corresponding DSS-colitis group in response to the lowest doses of *T. obliquus* (Mi175.B1.a) while a trend to significant decrease was described for microalgae treatment 100 mg/L (Figure 4.a.). This ratio was directly correlated to the severity of the colonic damage in this experimental model of colitis. Figure 4.b. showed that the preventive supplementation with *T. obliquus* (Mi175.B1.a) 100 mg/L significantly decreased colitis lesions. The evaluation of the weight of caecum demonstrated that DSS induced a reduction of caecal weight not counterbalanced by microalgae extract treatments.

**Figure 4.**
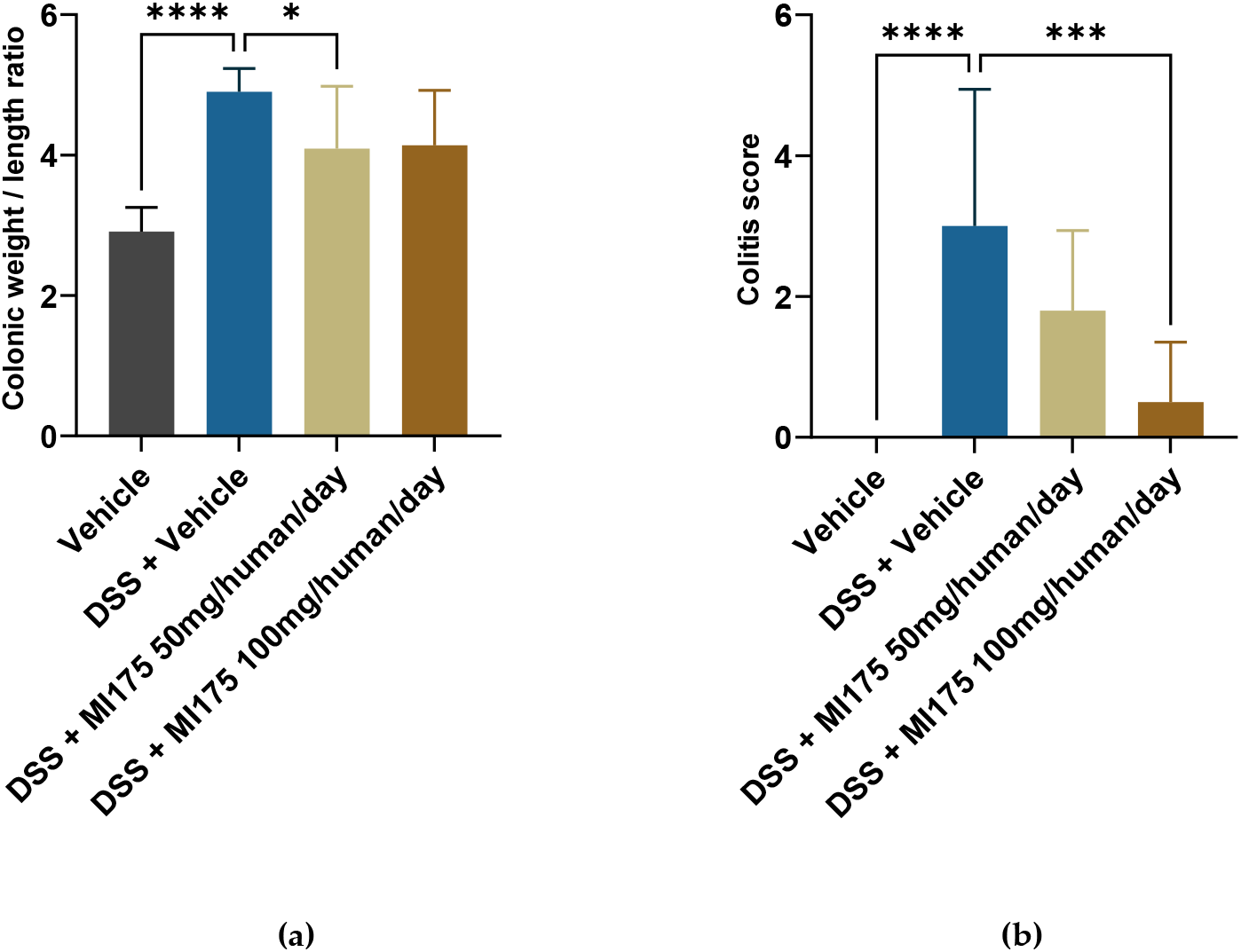
DSS-induced colitis macroscopic scoring. Effects of an oral administration of vehicle or 2 different doses of microalgal strain *T. obliquus* Mi175.B1.a extract on (a) colonic weight/length ratio and (b) colitis scores. * indicates significant difference between conditions using one-way ANOVA followed by Tukey’s post-hoc test. * means a p<0,05; ** means a p<0,01, *** means a p<0,001 and **** means a p<0,0001. n=9-10/ group.

**Figure 5.**
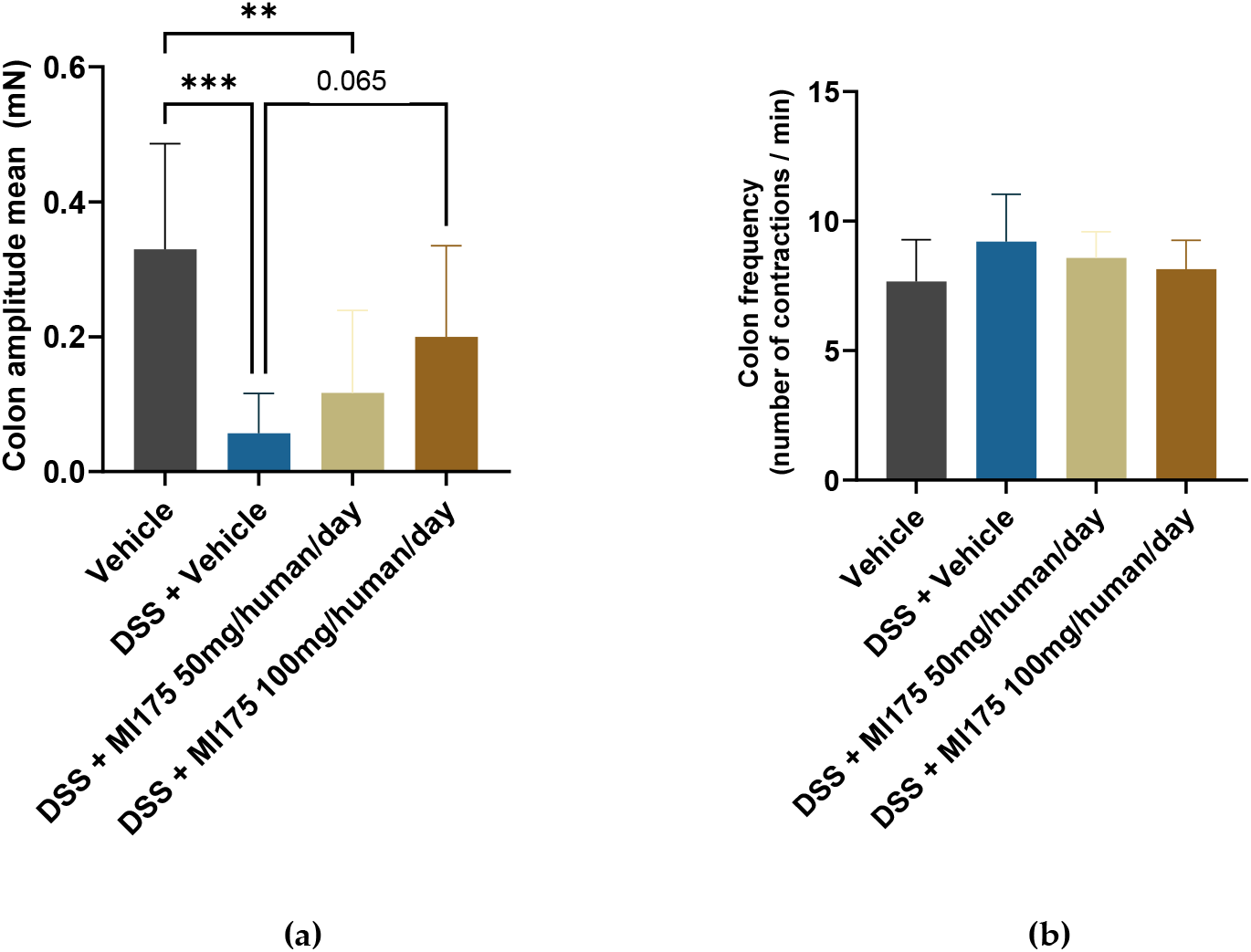
DSS-induced colitis macroscopic scoring. Effects of an oral administration of vehicle or 2 different doses of microalgal strain *T. obliquus* Mi175.B1.a extract on (a) colon amplitude and (b) colon frequency. * indicates significant difference between conditions using one-way ANOVA followed by Tukey’s post-hoc test. * means a p<0,05; ** means a p<0,01, *** means a p<0,001 and **** means a p<0,0001. n=9-10/ group.

#### 2.2.2. Colon motility assessment

The ulcerative colitis-like inflammation induced by DSS treatment altered the intrinsic sensory neurons excitability, disturbs enteric cholinergic neurotransmission, and suppresses smooth muscle reactivity. These effects lead to diarrhea induced by DSS treatment. The couple enteric nervous system / smooth muscle cells are major actors involved in the regulation of intestinal motility. In DSS mice, the daily oral administration of *T. obliquus* Mi175.B1.a extract 100 mg/L trend to significantly restore colon motility by increasing the amplitude of contraction compared to DSS-colitis group without impacting on the frequency of these contractions (Figure 4). Furthermore, we noted a beneficial effect of Mi175 100 mg/L but without reach the significance.

## 3. Discussion

Given the high prevalence of gastrointestinal (GI) disorders and symptoms, identifying effective therapeutics and preventive approaches to improve the gut microbiome and maintain intestinal barrier integrity is of critical importance. Microalgae have emerged as a promising source of natural nutraceuticals, which may play a critical role in maintaining the structural integrity of the gut mucosal barrier and modulating immune responses. In this study, our aim was to evaluate the efficacy of *Tetradesmus obliquus* Mi175.B1.a extract through two complementary approaches: (1) an in vitro model using the HT-29 cell line to assess cytotoxicity, intestinal permeability, immunomodulatory properties, and gut barrier function, and (2) an in vivo mouse model of dextran sodium sulfate (DSS)-induced colitis to examine parameters relevant to digestive physiology. The results provide novel and compelling evidence that *T. obliquus* extract enhances gut barrier integrity and modulates intestinal inflammation, as demonstrated through a dual *in vitro* and *in vivo* approach. The extract not only mitigated epithelial damage and inflammatory signaling in lipopolysaccharide (LPS)-challenged HT-29 cells but also significantly improved disease parameters and colon motility in a mouse model of dextran sodium sulfate (DSS)-induced colitis. These findings contribute to the growing recognition of microalgae as a promising source of gut-targeted nutraceuticals and reinforce the therapeutic potential of *T. obliquus* extract in intestinal health.

An essential aspect of evaluating the therapeutic potential of a novel nutraceutical is establishing its safety profile. In the present study, *Tetradesmus obliquus* Mi175.B1.a extract was administered orally to mice for a total duration of 21 days at two different concentrations. Throughout the treatment period, no signs of systemic toxicity or adverse clinical outcomes were observed. Specifically, body weight gain trajectories remained consistent with the control group, and no abnormalities in behavior, grooming, or food intake were detected. Moreover, macroscopic examination of the gastrointestinal tract showed no ulceration, bleeding, or signs of irritation in non-colitic animals receiving the extract. These findings suggest that *T. obliquus* extract is well tolerated even under inflammatory conditions, as it did not exacerbate the DSS-induced colitis model nor induce any additive stress. This safety profile is consistent with previous preclinical reports highlighting the low toxicity of microalgal species such as *Chlorella, Nannochloropsis*, and *Spirulina*, which are widely recognized as safe by regulatory bodies including the European Food Safety Authority (EFSA) and the U.S. FDA [41,42]. Although specific toxicological data for *T. obliquus* are still limited, its biochemical composition—characterized by natural pigments, essential fatty acids, and proteins devoid of known harmful secondary metabolites—supports its biocompatibility and was approuved as New Dietary Ingredient by FDA [43]. Importantly, the absence of cytotoxicity in HT-29 cells treated with both concentrations of *T. obliquus* extract over 72 hours further reinforces its safety for intestinal applications. These results, together with the tolerability observed *in vivo*, underscore the suitability of *T. obliquus* extract as a safe candidate for further development in the field of gut-targeted nutraceuticals.

Our *in vitro* findings indicate that *T. obliquus* Mi175.B1.a significantly reduced paracellular permeability, restored the expression of the tight junction protein Zonula occludens-1 (ZO-1), and downregulated pro-inflammatory markers such as TNF-α and Cox2. These observations align well with previous studies on other microalgae species. For instance, *Spirulina platensis* and *Chlorella vulgaris* have been shown to reinforce tight junction integrity and reduce inflammatory cytokine expression in intestinal epithelial cells under oxidative stress [44,45]. Similarly, PUFAs and carotenoids derived from microalgae like *Haematococcus pluvialis* and *Dunaliella salina* have been implicated in the regulation of intestinal inflammation through attenuation of the NF-κB and MAPK pathways [46].

Importantly, our *in vivo* results revealed that *T. obliquus* extract improved gut motility parameters, a critical component of digestive health. Proper colon motility is essential for maintaining bowel regularity, preventing microbial overgrowth, and supporting mucosal turnover and barrier function [47]. Impaired motility has been implicated in various disorders, including constipation, irritable bowel syndrome, and inflammatory bowel diseases [48]. The observed improvement in motility following *T. obliquus* supplementation is in line with previous reports on the motility-enhancing effects of dietary polyphenols, omega-3 fatty acids, and probiotics [21]. For example, Lactobacillus reuteri has been shown to accelerate colonic transit and reduce inflammation-induced dysmotility in murine colitis models [49], while plant-derived compounds like resveratrol and curcumin have demonstrated pro-motility and anti-inflammatory effects in experimental colitis [50]. Interestingly and as previously mentioned, the beneficial effects observed with *T. obliquus* extract are comparable to those described for well-characterized probiotic strains, such as Lactobacillus rhamnosus GG and Bifidobacterium longum, in similar models of gut barrier dysfunction and intestinal inflammation. In HT-29 cells, these probiotics have been shown to upregulate ZO-1 and occludin expression, decrease LPS-induced permeability, and suppress pro-inflammatory cytokine production via NF-κB inhibition [51,52]. Moreover, in DSS-induced colitis models, probiotics consistently reduce disease activity index (DAI), attenuate histological damage, and restore mucosal integrity through modulation of epithelial signaling pathways and immune responses [53]. The parallel between the outcomes of *T. obliquus* extract and probiotics suggests that certain microalgal bioactive may act through mechanisms akin to those employed by beneficial microbes—such as promoting tight junction integrity, modulating host immune responses, and possibly influencing gut microbiota composition. Therefore, *T. obliquus* may offer a promising alternative or complementary approach to probiotic therapy, especially in populations where probiotic viability or tolerance is a concern. The reproducibility of these effects across distinct therapeutic classes (microalgae and probiotics) further validates the robustness of the observed benefits and highlights the potential for *T. obliquus* extract to be incorporated into functional food or symbiotic formulations.

The observed biological activities of *T. obliquus* Mi175.B1.a extract are likely mediated by a synergistic network of bioactive compounds. This microalgae extract contains several interesting bioactive such as carotenoids (particularly lutein), polyunsaturated fatty acids (e.g., α-linolenic acid, EPA), phytosterols and antioxidant peptides [46,54–56]. Lutein, for example, has been shown to activate AMPK signaling, enhance the expression of tight junction proteins, and inhibit oxidative stress-induced damage to intestinal epithelial cells [57]. PUFAs in *T. obliquus* extract may also activate PPAR-γ and suppress pro-inflammatory cytokines through inhibition of NF-κB and MAPK pathways [54,58]. These mechanisms contribute to both the preservation of barrier integrity and the attenuation of inflammatory signaling. Moreover, the sulfated polysaccharides present in *T. obliquus* extract may act as prebiotic agents by modulating gut microbiota composition and enhancing the production of short-chain fatty acids (SCFAs), particularly butyrate—a critical metabolite known to reinforce tight junction function and promote mucosal healing [59,60]. These polysaccharides can also engage pattern recognition receptors such as toll-like receptors (TLRs), thereby downregulating inflammatory cascades and promoting immune tolerance [61]. Moreover, the anti-inflammatory activity of extracts from microalgae has already been demonstrated in other study models: in studies of age-related cognitive decline, an extract of the diatom *Phaeodactylum tricornutum* induced a reduction in Tnf-alpha and IL-6 in the hippocampus of mice, despite the induction of strong oxidative stress by injection of b-galactose, and the same extract induced a reduction in plasma CRP concentrations in an aged human population [62,63]. Additionally, microalgae-derived polyphenols and peptides have demonstrated the capacity to suppress TNF-α and IL-6 while upregulating mucins and barrier-forming proteins like occludin and claudins, further supporting the gut-protective profile of the extract [64]. Collectively, these multifaceted actions suggest that *T. obliquus* extract may exert its beneficial effects through a combination of antioxidant, anti-inflammatory, immune-modulatory, and microbiota-interactive pathways. Future studies employing transcriptomics, proteomics, and metabolomics will be essential to delineate the specific bioactive molecules responsible and to clarify their molecular targets in gut epithelium and immune cells.

Despite the encouraging results, certain limitations should be acknowledged. First, the study was performed exclusively on male mice, which does not account for potential sex-based differences in gut immune function, microbiota composition, or metabolite bioavailability. Future studies should include both sexes to improve translational relevance. Second, although the tested doses of *T. obliquus* Mi175.B1.a extract were well-tolerated over a 21-day period, the long-term safety and efficacy of chronic administration have yet to be determined. Additionally, the extract was evaluated as a whole, without isolating its individual bioactive constituents. While this holistic approach reflects real-world nutraceutical use, detailed compound profiling and fractionation are needed to identify the specific agents driving the observed benefits. Furthermore, this study did not assess microbiota composition, which is an important mediator of gut health and a potential mechanism of action for microalgal bioactive. Integration of metagenomic and metabolomic analyses in future experiments will help elucidate the interactions between *T. obliquus* Mi175.B1.a extract and the gut microbial ecosystem.

From a clinical translation perspective, well-designed human trials will be required to confirm efficacy and safety, highlighted in the present study, in target populations, such as individuals with inflammatory bowel disease, irritable bowel syndrome, or increased intestinal permeability. In this sense, a recently published double-blind randomized controlled trial showed that 4-week supplementation with *T. obliquus* strain Mi175.B1.a improves GI symptoms, potentially through effects on the gut microbiota, and may promote positive effects on mental health in healthy adults experiencing mild to moderate gastrointestinal GI distress [65]. Dose optimization, bioavailability studies, and delivery formulation (e.g., encapsulation) will also be crucial for successful nutraceutical development. The current findings position *Tetradesmus obliquus* Mi175.B1.a extract as a promising candidate for development into a functional food ingredient or dietary supplement aimed at enhancing gastrointestinal health. Its dual capacity to restore epithelial barrier function and reduce inflammatory responses supports its application in both preventive and therapeutic settings. Given the increasing recognition of intestinal permeability and chronic low-grade inflammation in systemic diseases—including metabolic syndrome, food allergies, and neuroinflammatory conditions—*T. obliquus* could also be explored in broader health contexts beyond traditional gastrointestinal disorders.

## 4. Materials and Methods

### 4.1. Microalgae extract production and characterization

The microalgae extract of the *Tetradesmus obliquus* Mi175.B1.a. production process was based on microalgae cultures in industrial photobioreactors, and all steps were realized in Microphyt facilities (Baillargues, FRANCE). Microphyt has developed and operates its own proprietary production technology called CAMARGUE®, which allows controlled microalgae biomass production. CAMARGUE® technology consists of a 1200 m serpentine glass piping circuit, with a 76 mm inside diameter and 4.5 mm wall thickness, folded horizontally in 24 straight lines with a vertical height of 3 m, 0.3 m wide, realizing a 50 m long tubular fence **[66]**.

*Tetradesmus obliquus* Mi175.B1.a. is grown in semi-continuous mode in the CAMARGUE® photobioreactor. The biomass is harvested when cell density is sufficient, and centrifuged to concentrate it before drying. The dry biomass is then ground and extracted with absolute ethanol. Following this extraction stage, the residual biomass is separated by filtration and addition of activated carbon. The vitro study was carried out on these intermediate extracts, to facilitate incubation on HT29 cells. The ethanol is then removed by vacuum evaporation in presence of MCT Coco oil to fluidize the final product. 0.5% vitapherol is added as a stabilizer. The global specifications of the microalgae extract of T*etradesmus obliquus* Mi175.B1.a. used in the *in vivo* study is presented in Table 2 below.

**Table 2.** *Tetradesmus obliquus* Mi175.B1.a composition

|  | Composition | Methods |
| --- | --- | --- |
| MCT oil based on coconut oil | 65-80% (w/w) |  |
| Mix of non-GMO tocopherols– E 306 | 0.45- 0.55% (w/w) |  |
| *Tetradasmus obliquus* Mi175.B1.a extract | 20-35% (w/w) |  |
| Carbohydrates | ≤ 20% (w/w) | Calculation |
| Proteins | ≤ 5% (w/w) | Kjeldahl (Titrimetry) |
| Vitamins (E, B4, B3...) | 0.1-0.2% (w/w) | HPLC |
| Total lipids | ≥ 80% (w/w) | Gravimetry |
| Total PUFAs | 3.5-11.5% (w/w) | GC-FID |
| Total MUFAs | 1-5.5% (w/w) | GC-FID |
| Total omega 3 | 2.4-6.2% (w/w) | GC-FID |
| Total omega 6 | 1.3-4.0 % (w/w) | GC-FID |
| Total phospholipids | 0.3-0.9 % (w/w) | HPLC |
| Total sterols | 0.3-0.7 % (w/w) | GC-FID |
| Free fatty acids | 7.6-20.6% (w/w) | GC-FID |
| Glycerides excluding MCT | 0.8-1.8%(w/w) | GC-FID |
| Lutein | 0.2 – 0.4% (w/w) | HPLC |
| Total carotenoids | 0.4-0.6 % (w/w) | Spectrophotometry |
| Chlorophylls | 0.06% (w/w) | Spectrophotometry |
| Humidity | ≤5% (w/w) | Thermogravimetry |

### 4.2. In vitro model methodology

This section was summarized in Figure 6.

**Figure 6.**
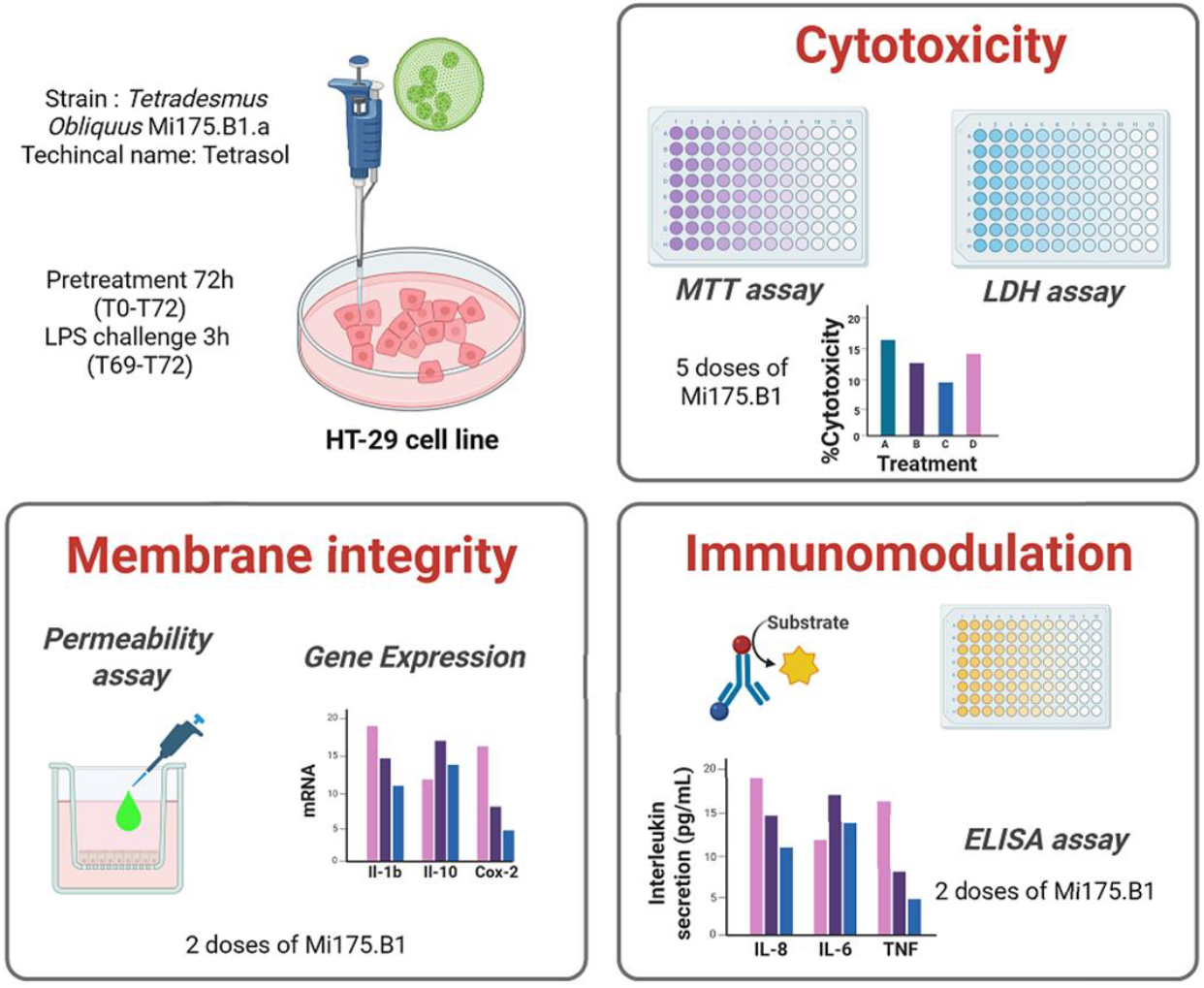
Study experimental design of *in vitro* model (created with Biorender). 4.2.1.Cell culture

#### 4.2.1. Cell culture

HT-29 cell line were cultured at 37°C in a humidified 5% CO_2_ incubator in Dul-becco modified eagle medium (DMEM) with high concentration of glucose, supplemented with 10% heat-inactivated fetal bovine serum (FBS), and 1% Penicillin-Streptomycin (P/S). The cell line adheres to the flasks and were therefore removed using trypsin/EDTA solution. The media was changed every day to maintain a high glucose concentration. Prior to the permeability experiments, HT-29 cells were seeded 0.4 μm, 1 cm^2^ transwell™ filters and grown to confluence. Then, cells were pretreated with two different concentrations of microalgal strain *T. obliquus* Mi175.B1.a (10 and 50 mg/L) for 72h following a stimulation with 1 µg/ml LPS for 3h. After the treatments, the supernatants and cell pellets were stored at -80°C until further analysis.

#### 4.2.2. Cell viability

The measurement of lactate dehydrogenase activity in the extracellular medium (LDH assay) associated with an indicator of cellular metabolic activity of cells (MTT assay) were used on 2D monolayer culture. To evaluate the toxicity of the tested items on IEC, the cells were treated with 5 concentrations of tested item (50, 10, 5, 1, 0.1 mg/L) for 24, 48 and 72h. Then, the supernatant was collected at the end of the treatment period. The activity of the lactate dehydrogenase released by damaged cells was quantified using the LDH assay. The viability of IEC was also evaluated by measuring the formazan produced by metabolic active cells using the MTT cell proliferation assay. The combination of these two cell viability tests allowed us to determine the two working concentrations of tested item.

#### 4.2.3. Gene expression

The RNA of the cells was extracted individually using TriReagent, according to the manufacturer’s instructions. Quantification analysis of total RNA was performed by analyzing 1 μl of each sample in Nanodrop. cDNA was prepared by reverse transcription of 500 ng total RNA using a RT iScript kit (Biorad). Real-time PCR was per-formed with the Via7 real-time PCR system and software (Applied biotechnology) using SYBR Green Real-Time PCR Master Mixes (Biorad) for detection, according to the manufacturer’s instructions. *Gapdh* was chosen as the housekeeping gene **[67]**. All samples were performed in duplicate, and data were analyzed according to the 2^ΔΔCT^ method. The identity and purity of the amplified product were assessed by melting curve analysis at the end of amplification. The IEC expression of the genes encoding for the following proteins were analyzed: zonula occludens 1 (*Zo-1*), claudin 4 (*Cldn4*), mucin 2 (*Muc2*), Interleukin-1β (*Il-1β*), Inter-leukin-10 (*Il-10*), Prostaglandin E2 (*Pge2*), Cyclooxygenase 2 (*Cox-2*), nuclear factor erythroid 2-related factor 2 (*Nrf-2*).

#### 4.2.4. Permeability assay

Paracellular permeability was assessed after treatment on another cell culture. The cells were gently rinsed with DPBS and incubated in the apical side with Hank’s balanced salt solution (HBSS) containing 1 mg/mL FITC-dextran (4.4 kDa) solution for 1 h. FITC-dextran flux was assessed by taking 100 μL from the basolateral chamber. Fluorescent signal was measured with a microplate reader using 492 nm excitation and 520 nm emission filters. FITC-dextran concentrations were determined using standard curves generated by serial dilutions. The concentration of FITC-dextran from the basolateral side represents the severity of permeability defect of HT-29 cells.

#### 4.2.5. Elisa assay

The supernatants were clarified by centrifugation (1000 g 20 min 4°C). The supernatants were stored at -80°C and were used for evaluating the immunomodulatory properties of the tested conditions using IL-6, IL-8, IL-10, IL-1β, TNFα ELISA kits (Merck, Germany). Measurements were assessed according to the manufacturer’s instructions.

### 4.3. In vivo model methodology

This section was summarized in Figure 7.

**Figure 7.**
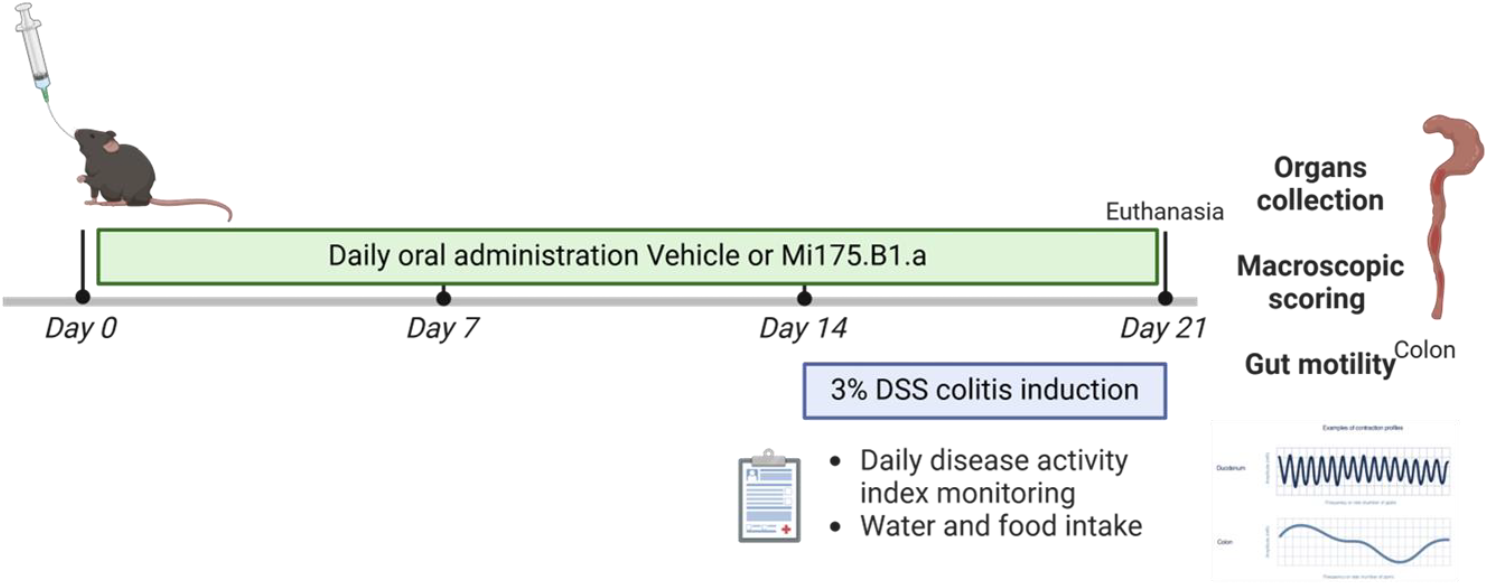
Study experimental design of an *in vivo* model (created with Biorender).

#### 4.3.1. Animals

Nine-week-old male C57BL/6J mice (Charles River Laboratory, l’Arbresle, France) were housed in specific pathogen free conditions and in a controlled environment (room temperature of 23 ± 2 °C, 12 h daylight cycle) with free access to food and water. The mice were randomly assigned to different groups of 10 animals. The non-colitic and the DSS-colitic groups received vehicle solution, whereas the microalgal strains groups were daily treated with the microalgae strain *T. Obliquus* Mi175.B1.a at the human equivalent doses of 50mg/day/70kg (Dose 1) and 100mg/day/70kg (Dose 2) for 21 days **[68,69]**. Two weeks after the onset of the study, colitis was induced in the DSS-colitis group by adding 3% DSS to drinking water for 6 days. The non-colitic group received normal tap water. Following euthanasia, 21 days after the onset of the study, the colon was collected, opened longitudinally, and all the contents collected; then it was weighed, and its length was measured. Immediately after euthanasia, the extent of colitis damage was assessed before tissue sampling. The macroscopic parameters analyzed were (ranging from 0 to 7): stool consistency (0 to 2 = formed to severe diarrhea), tissue adhesions (0 to 2 = absence to severe adhesions) and colonic damage (0 to 3 = normal to severe). A total score of 8 was assigned when animals died from colitis. All animal procedures were conducted in strict adherence to the European Union directive of 22 September 2010 (2010/63/UE). The experiment design (protocol APAFIS #49139-202404121614394 v6) was approved by the n°122 Ethics Committee in Toulouse.

#### 4.3.2. Daily disease activity index

The stool consistency and bleeding were evaluated daily for each mouse. These parameters were assigned a score according to the criteria proposed by Camuesco et al., which was used to calculate an average daily disease activity index (DAI) for each animal. DAI value was the combined scores of weight loss, stool consistency, and bleeding divided by 3 **[70]**.

#### 4.3.3. Isotonic contraction

After dissection, colonic segments were washed and incubated in oxygenated Krebs-Ringer solution for 30 min at 37°C, attached to the isotonic transducer (MLT7006 Isotonic Transducer, Hugo Basile, Comerio, Italy), and immersed in an organ bath of the same medium maintained at 37 °C. The load applied to the lever was 1g (10 mN). Isotonic contractions were recorded on Labchart software (AD Instruments) following the transducer displacement. After attaching the intestinal segments, contractions were recorded for 10 min. The basal contractions were presented as average of amplitude and frequency of contraction.

### 4.4. Data analysis

The data were expressed as the mean ± SEM. Differences between the experimental groups were assessed where appropriate using an unpaired Student’s, one- or two-way ANOVA, followed by post-hoc test. Data were analyzed using GraphPad Prism version 8.00 for Windows (GraphPad Software, San Diego, CA, USA). The results were considered statistically significant at p < 0.05.

## 5. Conclusions

In conclusion, *T. obliquus* Mi175.B1.a extract demonstrated promising gut-protective properties in both *in vitro* and *in vivo* inflammatory models, with potential implications for gut barrier integrity and inflammatory modulation. The findings support further research into its application as a nutraceutical for gut-related inflammatory conditions. Future studies should clarify its mechanisms of action, microbiome interactions, and clinical efficacy in demographics with inflammatory bowel disease, irritable bowel syndrome, or increased intestinal permeability.

## Author Contributions

Conceptualization, P.A., A.A. and R.P.; methodology, P.A., A.A. and R.P.; validation, J.M., P.A., A.A. and R.P..; formal analysis, A.A.; investigation, A.A.; data curation, A.A.; writing—original draft preparation, J.M. and A.A.; writing—review and editing, J.M., P.A., A.A. and R.P..; supervision, P.A., A.A. and R.P. All authors have read and agreed to the published version of the manuscript.

## Institutional Review Board Statement

All animal procedures were conducted in strict adherence to the European Union di-rective of 22 September 2010 (2010/63/UE). The experiment design (protocol APAFIS #49139-202404121614394 v6) was approved by the n°122 Ethics Committee in Toulouse.

## Data Availability Statement

Raw data are available from corresponding author upon justified requests.

